# Algorithmic methods to infer the evolutionary trajectories in cancer progression

**DOI:** 10.1101/027359

**Authors:** Giulio Caravagna, Alex Graudenzi, Daniele Ramazzotti, Rebeca Sanz-Pamplona, Luca De Sano, Giancarlo Mauri, Victor Moreno, Marco Antoniotti, Bud Mishra

## Abstract

The genomic evolution inherent to cancer relates directly to a renewed focus on the voluminous next generation sequencing (NGS) data, and machine learning for the inference of explanatory models of how the (epi)genomic events are choreographed in cancer initiation and development. However, despite the increasing availability of multiple additional - omics data, this quest has been frustrated by various theoretical and technical hurdles, mostly stemming from the dramatic heterogeneity of the disease. In this paper, we build on our recent works on “selective advantage” relation among driver mutations in cancer progression and investigate its applicability to the modeling problem at the population level. Here, we introduce PiCnIc (Pipeline for Cancer Inference), a versatile, modular and customizable pipeline to extract ensemble-level progression models from cross-sectional sequenced cancer genomes. The pipeline has many translational implications as it combines state-of-the-art techniques for sample stratification, driver selection, identification of fitness-equivalent exclusive alterations and progression model inference. We demonstrate PiCnIc’s ability to reproduce much of the current knowledge on colorectal cancer progression, as well as to suggest novel experimentally verifiable hypotheses.

Statement of Significance: *A causality based new machine learning Pipeline for Cancer Inference* (PicNic) *is introduced to infer the underlying somatic evolution of ensembles of tumors from next generation sequencing data*. PicNic *combines techniques for sample stratification, driver selection and identification of fitness-equivalent exclusive alterations to exploit a novel algorithm based on Suppes’ probabilistic causation. The accuracy and translational significance of the results are studied in details, with an application to colorectal cancer*. PicNic *pipeline has been made publicly accessible for reproducibility, interoperability and for future enhancements*.

## 1 Introduction

Since the late seventies evolutionary dynamics, with its interplay between variation and selection, has progressively provided the widely-accepted paradigm for the interpretation of cancer emergence and development [1–3]. Random alterations of an organism’s (epi)genome can sometimes confer a functional *selective advantage*^1^ to certain cells, in terms of adaptability and ability to survive and proliferate. Since the consequent *clonal expansions* are naturally constrained by the availability of resources (metabolites, oxygen, etc.), further mutations in the emerging heterogeneous tumor populations are necessary to provide additional *fitness* of different kinds that allow survival and proliferation in the unstable micro environment. Such further advantageous mutations will eventually allow some of their sub-clones to outgrow the competing cells, thus enhancing tumor’s heterogeneity as well as its ability to overcome future limitations imposed by the rapidly exhausting resources. Competition, predation, parasitism and cooperation have been in fact theorized as co-present among cancer clones [4].

In the well-known vision of Hanahan and Weinberg [5,6], the phenotypic stages that characterize this multistep evolutionary process are called *hallmarks*. These can be acquired by cancer cells in many possible alternative ways, as a result of a complex biological interplay at several spatio-temporal scales that is still only partially deciphered [7]. In this framework, we distinguish “alterations” driving the hallmark acquisition process (i.e., *drivers)* by activating *oncogenes* or inactivating *tumor suppressor genes*, from those that are transferred to sub-clones without increasing their fitness (i.e., *passengers)* [8]. Driver identification is a modern challenge of cancer biology, as distinct cancer types exhibit very different combinations of drivers, some cancers display mutations in hundreds of genes [9], and the majority of drivers is mutated at low frequencies (“long tail” distribution), hindering their detection only from the statistics of the recurrence at the population-level [10].

Cancer clones harbour distinct types of alterations. The *somatic* (or *genetic)* ones involve either few nucleotides or larger chromosomal regions. They are usually catalogued as *mutations* - i.e., single nucleotide or structural variants at multiple scales (insertions, deletions, inversions, translocations) – of which only some are detectable as *Copy Number Alterations* (CNAs), most prevalent in many tumor types [11]. Also *epigenetic* alterations, such as *DNA methylation* and *chromatin reorganization*, play a key role in the process [12]. The overall picture is confounded by factors such as *genetic instability* [13], *tumor-microenvironment* interplay [14,15], and by the influence of *spatial organization* and *tissue specificity* on tumor development [16]^2^.

Significantly, in many cases, distinct driver alterations can damage in a similar way the same *functional pathway*, leading to the acquisition of new hallmarks [17–21]. Such alterations individually provide an equivalent *fitness gain* to cancer cells, as any additional alteration hitting the same pathway would provide no further selective advantage. This dynamic results in groups of driver alterations that form *mutually exclusive* patterns across tumor samples from different patients (i.e., the sets of alterations that are involved in the same pathways tend not to occur mutated together). This phenomenon has significant translational consequences.

An immediate challenge posed by this state of affairs is the dramatic *heterogeneity* of cancer, both at the *inter-tumor* and at the *intra-tumor* levels [22]. The former manifests as different patients with the same cancer type can display few common alterations. This obsersvation led to the development of techniques to stratify tumors into *subtypes* with different genomic signatures, prognoses and response to therapy [23]. The latter form of heterogeneity refers to the observed genotypic and phenotypic variability among the cancer cells within a single neoplastic lesion, characterized by the coexistence of more than one cancer clones with distinct evolutionary histories [24].

Cancer heterogeneity poses a serious problem from the diagnostic and therapeutic perspective as, for instance, it is now acknowledged that a single biopsy might not be representative of other parts of the tumor, hindering the problem of devising effective treatment strategies [4]. Therefore, presently the quest for an extensive etiology of cancer heterogeneity and for the identification of cancer evolutionary trajectories is central to cancer research, which attempts to exploit the massive amount of sequencing data available through public projects such as The Cancer Genome Atlas (TCGA) [25].

Such projects involve an increasing number of *cross-sectional* (epi)genomic profiles collected via single biopsies of patients with various cancer types, which might be used to extract trends of cancer evolution across a population of samples^3^. Higher resolution data such as *multiple samples* collected from the same tumor [24], as well as *single-cell* sequencing data [26], might be complementarily used to face the same problem within a specific patient. However, the lack of public data coupled to the problems of accuracy and reliability, currently prevents a straightforward application [27].

These different perspectives lead to the different mathematical formulations of the problem of *inferring a cancer progression model* from genomic data, and a need for versatile computational tools to analyze data reproducibly – two intertwined issues examined at length in this paper [28]. Indeed, such models and tools can be focused either on characteristics of a population, i.e. *ensemble-level*, or on multiple clonality in a *single-patient*. In general, both problems deal with understanding the *temporal ordering of somatic alterations* accumulating during cancer evolution, but use orthogonal perspectives and different input data – see Figure 1 for a comparison. This paper proposes a new computational approach to efficiently deal with various aspects of the problem at a patient population level, relegating the other aspects to future publications.

**Figure 1:**
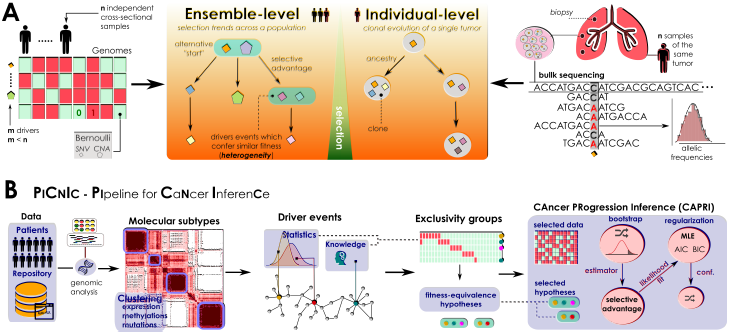
A. Problem statement. (left) Inference of ensemble-level cancer progression models from a cohort of *n* independent patients (cross-sectional). By examining a list of somatic mutations or CNAs per patient (0/1 variables) we infer a probabilistic graphical model of the temporal ordering of fixation and accumulation of such alterations in the input cohort. Sample size and tumor heterogeneity complicate the problem of extracting population-level trends, as this requires accounting for patients’ specificities such as multiple starting events. (right) For an individual tumor, its clonal phylogeny and prevalence is usually inferred from multiple biopsies or single-cell sequencing data. Phylogeny-tree reconstruction from an underlying statistical model of reads coverage or depths estimates alterations’ prevalence in each clone, as well as ancestry relations. This problem is mostly worsened by the high intra-tumor heterogeneity and sequencing issues. **B.** The PicNic pipeline for ensemble-level inference includes several sequential steps to reduce tumor heterogeneity, before applying the CAPRI [40] algorithm. Available mutation, expression or methylation data are first used to stratify patients into distinct tumor molecular subtypes, usually by exploiting clustering tools. Then, subtype-specific alterations driving cancer initiation and progression are identified with statistical tools and on the basis of prior knowledge. Next is the identification of the fitness-equivalent groups of mutually exclusive alterations across the input population, again done with computational tools or biological priors. Finally, CAPRI processes a set of relevant alterations within such groups. Via bootstrap and hypothesis-testing, CAPRI extracts a set of “selective advantage relations” among them, which is eventually narrowed down via maximum likelihood estimation with regularization (with various scores). The ensemble-level progression model is obtained by combining such relations in a graph, and its confidence is assessed via various bootstrap and cross-validation techniques.

**Ensemble-level cancer evolution.** It is thus desirable to extract a *probabilistic graphical model* explaining the statistical trend of accumulation of somatic alterations in a population of *n* crosssectional samples collected from patients diagnosed with a specific cancer. To normalize against the experimental conditions in which tumors are sampled, we only consider the *list of alterations detected per sample* – thus, as 0/1 Bernoulli random variables.

Much of the difficulty lies in estimating the true and unknown trends of *selective advantage* among genomic alterations in the data, from such observations. This hurdle is not unsurmountable, if we constrain the scope to only those alterations that are *persistent across tumor evolution in all sub-clonal populations*, since it yields a consistent model of a temporal ordering of mutations. Therefore, epigenetic and trascriptomic states, such as hyper and hypo-methylations or over and under expression, could only be used, provided that they are persistent through tumor development [29].

Historically, the linear model of colorectal tumor progression by Vogelstein is an instance of an early solution to the cancer progression problem [30]. That approach was later generalized to accommodate *tree-models of branched evolution* [31–34] and later, further generalized to the inference of *directed acyclic graph* models, with several distinct strategies [35–38]. We contributed to this research program with the `Cancer Progression Extraction with Single Edges (CAPRESE)` and the `Cancer Progression Inference (CAPRI)` algorithms, which are currently implemented in `TRONCO`, an open source `R` package for `Translational Oncology` available in standard repositories [39–41]. Both techniques rely on Suppes’ theory of probabilistic causation to define estimators of selective advantage [42], are robust to the presence of noise in the data and perform well even with limited sample sizes. The former algorithm exploits shrinkage-like statistics to extract a tree model of progression, the latter combines bootstrap and maximum likelihood estimation with regularization to extract general directed acyclic graphs that capture branched, independent and confluent evolution. Both algorithms represent the current state-of-the-art approach to this problem, as they outperform others in speed, scale and predictive accuracy.

**Clonal architecture in individual patients.** A closely related problem addresses the detection of clonal signatures and their prevalence in individual tumors, a problem complicated by *intra-tumor* heterogeneity.

Even though this phylogenetic version of the progression inference problem naturally relies on data produced from *single-cell sequencing* assays [43,44], the majority of approaches still make use of *bulk sequencing* data, usually from multiple biopsies of the same tumors [24,45]. Indeed, several approaches try to extract the clonal signature of single tumors from *allelic imbalance proportions*, a problem made difficult as sequenced samples usually contain a large number of cells belonging to a collection of sub-clones resulting from the complex evolutionary history of the tumor [46–55].

We keep the current work focused on the inference of progression models at the ensemble level, and plan to return to this variant to the problem in another publication.

## 2 The PicNic pipeline

We report on the design, development and evaluation of the Pipeline for Cancer Inference (PicNic) to extract ensemble-level cancer progression models from cross-sectional data (Figure 1). PicNic is versatile, modular and customizable; it exploits state-of-the-art data processing and machine learning tools to:

1. identify *tumor subtypes* and then in each subtype;
2. select (epi)genomic events *relevant* to the progression;
3. identify groups of events that are likely to be observed as *mutually exclusive*;
4. infer *progression models* from groups and related data, and annotate them with associated statistical confidence.

All these steps are necessary to minimize the confounding effects of inter-tumor heterogeneity, which are likely to lead to wrong results when data is not appropriately pre-processed^4^.

In each stage of `PicNic` different techniques can be employed, alternatively or jointly, according to specific research goals, input data, and cancer type. Prior knowledge can be easily accommodated into our pipeline, as well as the computational tools discussed in the next subsections and summarized in Figure 2. The rationale is similar in spirit to workflows implemented by consortia such as `TCGA` to analyze huge populations of cancer samples [56,57]. One of the main novelties of our approach, is the exploitation of groups of exclusive alterations as a proxy to detect fitness-equivalent trajectories of cancer progression. This strategy is only feasible by the hypothesis-testing features of the recently developed `CAPRI` algorithm, an algorithm uniquely addressing this crucial aspect of the ensemble-level progression inference problem [40].

**Figure 2:**
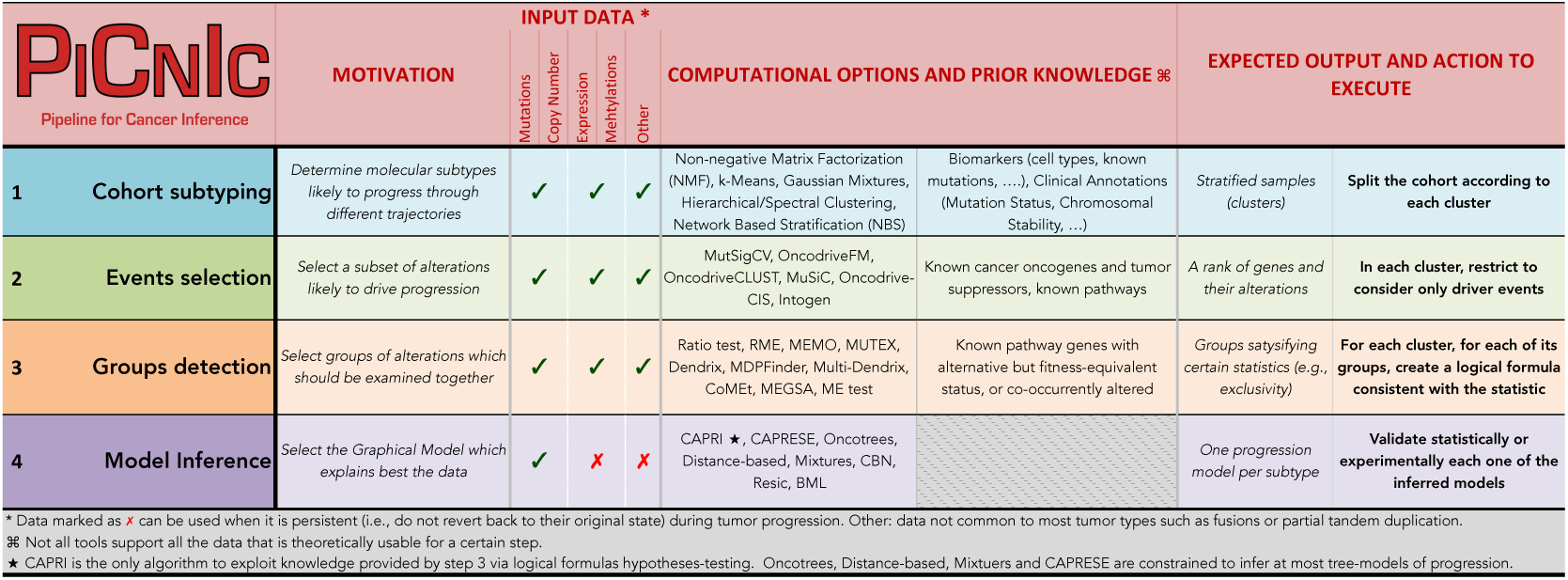
The PicNic pipeline. We do not provide a unique all-encompassing rationale to instantiate PicNic as all steps refer to research area currently development, where the optimal approach is often dependent on the type of data available and prior knowledge about the cancer under study. References are provided for each tool that can be used to instantiate PicNic: NMF [61], k-Means, Gaussian Mixtures, Hierarchical/Spectral Clustering [62], NBS [66], MutSigCV [68], OncodriveFM [69], OncodriveCLUST [70], MuSiC [71] Oncodrive-CIS [72] Intogen [73], Ratio [74], RME [75], MEMO [76], MUTEX [77], Dendrix [78], MDPFinder [79], Multi-Dendrix [80], CoMEt [81], MEGSA [82], ME [83], CAPRI [40], CAPRESE [39], Oncotrees [31, 33], Distance-based [32], Mixtures [34], CBN [35, 36], Resic [37] and BML [38].

In the Results section, we study in details a specific use-case for the pipeline, processing colorectal cancer data from TCGA, where it is able to re-discover much of the existing body of knowledge about colorectal cancer progression. Based on the output of this pipeline, we also propose novel experimentally-verifiable hypotheses.

### 2.1 Reducing inter-tumor heterogeneity by cohort subtyping

In general, for each of *n* tumors (patients) we assume relevant (epi)genetic data to be available. We do not put constraints on data gathering and selection, leaving the user to decide the appropriate “resolution” of the input data. For instance, one might decide whether somatic mutations should be classified by type or by location, or aggregated. Or, one might decide to lift focal CNAs to the lower resolution of cytobands or full arms (e.g., in a kidney cancer cohort where very long CNAs are more common than focal events [58]). These choices depend on data and on the overall understanding of such alterations and their functional effects for the cancer under study, and no single all-encompassing rationale may be provided.

With these data at hand, we might wish to identify cancer subtypes in the *heterogeneous mixture* of input samples. In some cases the classification can benefit from clinical biomarkers, such as evidences of certain cell types [59], but in most cases we will have to rely on multiple *clustering* techniques at once, see, e.g., [56,57]. Many common approaches cluster expression profiles [60], often relying on non-negative matrix factorization techniques [61] or earlier approaches such as k-means, Gaussians mixtures or hierarchical/spectral clustering - see the review in [62]. For glioblastoma and breast cancer, for instance, mRNA expression subtypes provides good correlation with clinical phenotypes [63–65]. However, this is not always the case as, e.g., in colorectal cancer such clusters mismatch with survival and chemotherapy response [63]. Clustering of *full exome* mutation profiles or smaller panels of genes might be an alternative as it was shown for ovarian, uterine and lung cancers [66,67].

Using pipelines such as PicNic, we expect that the resulting subtypes will be routinely investigated, eventually leading to distinct progression models, which shall be characteristic of the population-level trends of cancer initiation and progression.

### 2.2 Selection of driver events

In subtypes detection, it becomes easier to find similarities across input samples when more alterations are available, as features selection gains precision. In progression inference, instead, one wishes to focus on *m* ≪ *n driver* alterations, which ensure also an appropriate statistical ratio between sample size (*n*, here the subtype size) and problem dimension (*m*).

Multiple tools filter out driver from passenger mutations. `MutSigCV` identifies drivers mutated more frequently than background mutation rate [68]. `OncodriveFM` avoids such estimation but looks for functional mutations [69]. `OncodriveCLUST` scans mutations clustering in small regions of the protein sequence [70]. `MuSiC` uses multiple types of clinical data to establish correlations among mutation sites, genes and pathways [71]. Some other tools search for driver CNAs that affect protein expression [72]. All these approaches use different statistical measures to estimate signs of positive selection, and we suggest using them in an orchestrated way, as done by platforms such as `Intogen` [73].

We anticipate that such tools will run independently on each subtype, as driver genes will likely differ across them, mimicking the different molecular properties of each group of samples; also, lists of genes produced by these tools might be augmented with prior knowledge about tumor suppressors or oncogenes.

### 2.3 Fitness equivalence of exclusive alterations

When working at the ensemble-level, identification of “groups of mutually exclusive” alterations is crucial to derive a correct inference. This step of `PicNic` is another attempt to resolve part of the inter-tumor heterogeneity, as such alterations *could* lead to the same phenotype (i.e., hence resulting “equivalent” in terms of progression), despite being genotypically “alternative”, i.e., exclusive, across the input cohort. This information shall be used to detect alternative routes to cancer progression which capture the specificities of individual patients.

A plethora of recent tools can be used to detect groups of fitness equivalent alterations, according to the data available for each subtype; greedy approaches [74, 75] or their optimizations, such as `MEMO`, which constrain search-space with network priors [76]. This strategy is further improved in `MUTEX`, which scans mutations and focal CNAs for genes with a common downstream effect in a curated signalling network, and selects only those genes that significantly contributes to the exclusivity pattern [77]. Other tools such as `Dendrix`, `MDPFinder`, `Multi-Dendrix`, `CoMEt`, `MEGSA` or `ME`, employ advanced statistics or generative approaches without priors [78–83].

In such groups, we distinguish between *hard* and *soft* forms of exclusivity, the former assuming strict exclusivity among alterations, with random errors accounting for possible overlaps (i.e., the majority of samples do not share alterations from such groups), the latter admitting co-occurrences (i.e., some samples might have common alterations, within a group) [77].

`CAPRI` is currently the only algorithm which incorporates this type of information, in inferring a model. Each of these groups are in fact associated with a *“testable hypothesis”* written in the well-known language of *propositional Boolean formulas*^5^. Consider the following example: we might be informed that APC and Ctnnb1 *mutations* show a trend of soft-exclusivity in our cohort - i.e., some samples harbor both mutations, but the majority just one of the two mutated genes. Since such mutations lead to *β*-catenin deregulation (the phenotype), we might wonder whether such state of affairs could be responsible for progression initiation in the tumors under study. An affermative response would equate, in terms of progression, the two mutations. To *test this hypothesis*, one may spell out formula apc ∨ ctnnb1 to `CAPRI`, which means that we are suggesting to the inference engine that, besides the possible evolutionary trajectories that might be inferred by looking at the two mutations as *independent*, trajectories involving such a “composite” event, shall be considered as well. It is then up to `CAPRI` to decide which, of all such trajectories, is significant, in a *statistical sense*.

In general, formulas allow users to test general hypotheses about complex model structures involving multiple genes and alterations. These are useful in many cases: for instance, where we are processing samples which harbour homozygous losses or inactivating mutations in certain genes (i.e., equally disruptive genomic events), or when we know in advance that certain genes are controlling the same pathway, and we might speculate that a single hit in one of those decreases the selection pressure on the others. We note that, with no hypothesis, a model with such alternative trajectories *cannot be analyzed, due to various computational* limitations inherent to the inferential algorithms (see [40]).

From a practical point of view, `CAPRI`’s formulas/hypotheses-testing features “help” the inference process, but do not “force” it to select a specific model, i.e., the *inference is not biased*. In this sense, the trajectories inferred by examining these composite model structures (i.e., the formulas) *are not given any statistical advantage* for inclusion in the final model. However, in spite of a natural temptation to generate as many hypotheses as possible, it is prudent to always limit the number of hypotheses according to the number of samples and alterations. Note that this approach can also be extended to accommodate, for instance, co-occurrent alterations in significantly mutated subnetworks [84,85].

### 2.4 Progression inference and confidence estimation

We use `CAPRI` to reconstruct cancer progression models of each identified molecular subtype, provided that there exist a reasonable list of driver events and the groups of fitness-equivalent exclusive alterations. Since currently `CAPRI` represents the state of the art, and supports complex formulas for groups of alterations detected in the earlier `PicNic` step, it was well-suited for the task.

`CAPRI`’s input is a binary *n* × (*m* * *k*) matrix **M** with *n* samples (a subtype size), *m* driver alteration events (0/1 Bernoulli random variables) and *k* testable formulas. Each sample in **M** is described by a binary sequence: the 1’s denote the presence of alterations. `CAPRI` first performs a computationally fast *scan* of **M** to identify a set ***S*** of plausible selective advantage relations among the driver alterations and the formulas; then, it reduces ***S*** to the most relevant ones, ***S***̂ ⊂ ***S*** Each relation is represented as an edge connecting drivers/formulas in a Graphical Model – which shall be termed Suppes-Bayes Causal Network. This network represents the *joint probability distribution*^6^ of observing a set of driver alterations in a cancer genome, subject to constraints imposed by Suppes’ *probabilistic causation* formalism [42].

Set ***S*** is built by a statistical procedure. Among any pair of input drivers/formulas *x* and *y*, `CAPRI` postulates that *x* → *y* ∈ ***S*** could be a selective advantage relation with “*x* selecting for *y*” if it estimates that two conditions hold

1. “ *x* is earlier than *y*”;
2. “*x*’s presence increases the probability of observing *y*”.

Such claims, grounded in Suppes’ theory of probabilistic causation, are expressed as inequalities over *marginal* and *conditional* distributions of *x* and *y*. These are assessed via a standard Mann-Withney U test after the distributions are estimated from a reasonable number (e.g., 100) of *non-parametric bootstrap resamples* of **M** (see Supplementary Material). `CAPRI`’s increased performance over existing methods can be motivated by the reduction of the state space within which models are searched, via ***S***.

Optimization of ***S*** is central to our tolerance to *false positives* and *negatives* in ****S***̂*. We would like to select only the minimum number of relations which are true and statistically supported, and build our model from those. `CAPRI`’s implementation in `TRONCO` [41] selects a subset by optimizing a *score function* which assigns to a model a real number equal to its *log-likelihood* (probability of generating data for the model) minus a *penalty term* for model complexity - a regularization term increasing with ***S***̂’s size, and hence penalizing overly complex models. It is a standard approach to avoid overfitting, and usually relies on the `Akaike` or the `Bayesian Information Criterion` (`AIC` or `BIC`) as regularizers. Both scores are approximately correct; `AIC` is more prone to overfitting but likely to provide also good predictions from data and is better when false negatives are more misleading than positive ones. `BIC` is more prone to underfitting errors, thus more parsimonious and better in opposite direction. As often done, we suggest approaches that to combine but distinguish which relations are selected by `BIC` versus `AIC`. Details on the algorithm are provided as see Supplementary Material.

**Statistical confidence of a model**. In-vitro and in-vivo experiments provide the most convincing validation for the newly suggested selective advantage relations and hypotheses, yet this is out of reach in some cases.

Nonetheless, statistical validation approaches can be used almost universally to assess the confidence of edges, parent sets and whole models, either via *hypothesis-testing* or *bootstrap* and *crossvalidation* scores for Graphical Models. We briefly discuss approaches that are implemented in `TRONCO`, and refer to the Supplementary Materials for additional details.

First, `CAPRI` builds ***S*** by computing two p-values per edge, for the confidence in condition (1) and (2). In addition, for each edge *x* → *y*, it computes a third p-value via hypergeometric testing against the hypothesis that the co-occurrence of *x* and *y* is due to chance. These p-values measure confidence in the direction of each edge and the amount of statistical dependence among *x* and *y*.

Second, for each model inferred with `CAPRI` we can estimate *(a posteriori*) how frequently our edges would be retrieved if we resample from our data *(non-parametric* bootstrap), or from the model itself, assuming its correctness *(parametric* bootstrap) [86]. Also, we can measure the bias in `CAPRI`’s construction of ***S*** due to the random procedure which estimates the distributions in condition (1) and (2) *(statistical* bootstrap).

Third, scores can be computed to quantify the consistency for the model against bias in the data and models. For instance, *non-exhaustive k*-fold cross-validation can be used to compute the *entropy loss* for the whole model, and the *prediction* and *posterior classification errors* for each edge or parent set [87].

## 3 Results

### 3.1 Evolution in a population of MSI/MSS colorectal tumors

It is common knowledge that *colorectal cancer* (CRC) is a heterogeneous disease comprising different molecular entities. Indeed, it is currently accepted that colon tumors can be classified according to their global genomic status into two main types: *microsatellite unstable tumors* (MSI), further classified as high or low, and *microsatellite stable* (MSS) tumors (also known as tumors with *chromosomal instability*). This taxonomy plays a significant role in determining pathologic, clinical and biological characteristics of CRC tumors [88]. Regarding molecular progression, it is also well established that each subtype arises from a distinctive molecular mechanism. While MSS tumors generally follow the classical adenoma-to-carcinoma progression described in the seminal work by Vogelstein and Fearon [89], MSI tumors result from the inactivation of DNA mismatch repair genes like mlh-1 [90].

With the aid of the `TRONCO` package, we instantiated `PicNic` to process colorectal tumors freely available through `TCGA` project `COADREAD` [56] (see Supplementary Figure S1), and inferred models for the MSS and MSI-HIGH tumor subtypes (shortly denoted MSI) annotated by the consortium. In doing so, we used a combination of background knowledge produced by `TCGA` and new computational predictions; to a different degree, some knowledge comes from manual curation of data and other from tools mentioned in `PicNic`’s description (see Figure 2). Data and exclusivity groups for MSI tumors are shown in Figure 3, the analogous for MSS tumors is provided as Supplementary Material.

**Figure 3:**
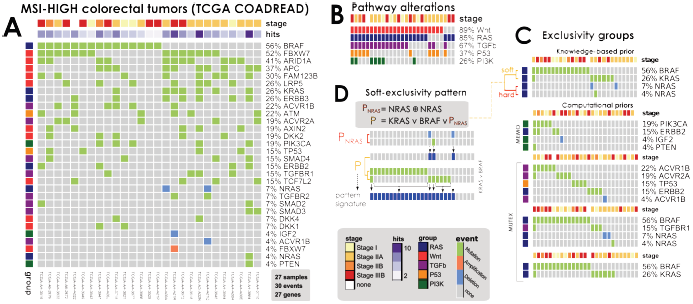
**A.** MSI-HIGH colorectal tumors from the TCGA COADREAD project [56], restricted to 27 samples with both somatic mutations and high-resolution CNA data available and a selection out of 33 driver genes annotated to wnt, ras, pi3k, tgf-*β* and p53 pathways. This dataset is used to infer the model in Figure 5. **B.** Mutations and CNAs in MSI-HIGH tumors mapped to pathways confirm heterogeneity even at the pathway-level. **C.** Groups of mutually exclusive alterations were obtained from [56] - which run the MEMO [76] tool - and by MUTEX [77] tool. In addition, previous knowledge about exclusivity among genes in the RAS pathway was exploited. **D.** A Boolean formula input to CAPRI tests the hypothesis that alterations in the RAS genes kras, NRAS and braf confer equivalent selective advantage. The formula accounts for hard exclusivity of alterations in NRAS mutations and deletions, jointly with soft exclusivity with kras and NRAS alterations.

For the models inferred, which are shown in Figures 4 and 5, we evaluated various forms of statistical confidence measured as p-values, bootstrap scores (in what follows, `npb` denotes non-parametric bootstrap and the closer to 100 the better), and cross-validation statistics reported in the Supplementary Material. Many of the postulated selective advantage relations (i.e., model edges) have very strong statistical support for COADREAD samples, although events with similar marginal frequency may lead to ambiguous imputed temporal ordering (i.e., the edge direction). In general, we observed that overall the estimates are slightly better in the MSS cohort (entropy loss < 1% versus 3.8%), which is expected given the difference in sample size of the two datasets (152 versus 27 samples), see Material and Methods for details.

**Figure 4:**
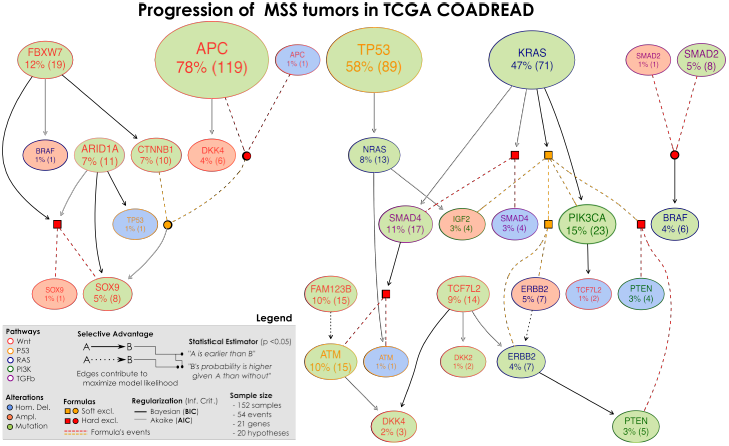
Selective advantage relations inferred by CAPRI constitute MSS progression; input dataset in Supplementary Figure S3 and S4. Formulas written on groups of exclusive alterations, e.g., sox9 amplifications and mutations, are displayed in expanded form; their events are connected by dashed lines with colors representing the type of exclusivity (red for hard, orange for soft), logical connectives are squared when the formula is selected, and circular when the formula selects for a downstream node. For this model of MSS tumors in COADREAD, we find strong statistical support for many edges (p-values, bootstrap scores and cross-validation statistics shown as Supplementary Material), as well as the overall model. This model captures both current knowledge about CRC progression - e.g, selection of alterations in pi3k genes by the KRAs mutations (directed or via the MEMO group, with BIC) – as well as novel interesting testable hypotheses – e.g., selection of sox9 alterations by fbxwT mutations (with BIC).

**Figure 5:**
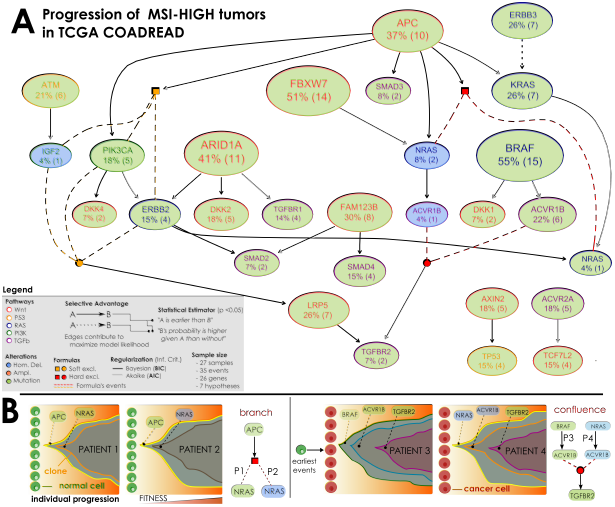
**A.** Selective advantage relations inferred by CAPRI constitute MSI-HIGH progression; input dataset in Figure 3. Formulas written on groups of exclusive alterations are expanded as in Figure 4. For each relation, confidence is estimated as for MSS tumors and reported as Supplementary Material. In general, this model is supported by weaker statistics than MSS tumors - possibly because of this small sample size (*n*=27). Still, we can find interesting relations involving apc mutations which select for pik3ca ones (via BIC) as well as selection of the MEMO group (erbb2/pik3ca mutations or iGf2 deletions) predicted by AIC. Similarly, we find a strong selection trend among mutations in eRbb2 and kras, despite in this case the temporal precedence among those mutations is not disentangled as the two events have the same marginal frequencies (26%). **B**. Evolutionary trajectories of clonal expansion predicted from two selective advantage relations in the model. apc-mutated clones shall enjoy expansion, up to acquisition of further selective advantage via mutations or homozygous deletions in NRAS. These cases should be representative of different individuals in the population, and the ensemble-level interpretation should be that “APC mutations select for NRAS alterations, in hard exclusivity” as no sample harbour both alterations. A similar argument can show that the clones of patients harbouring distinct alterations in acvr1b – and different upstream events – will enjoy further selective advantage from mutation in the tgfbr2 gene.

**Interpretation of the models.** Our models capture the well-known features distinguishing MSS and MSI tumors: for the former APC, KRAS and tf53 mutations as primary events together with chromosomal aberrations, for the latter braf mutations and lack of chromosomal alterations. Of all 33 driver genes, 15 are common to both models - e.g., apc, braf, kras, nras, tp53 and fam123b among others (mapped to pathways like wnt, mapk, apoptosis or activation of T-cell lymphocites), although in different relationships (position in the model), whereas new (previously un-implicated) genes stood out from our analysis and deserve further research.

*MSS (Microsatellite Stable)*. In agreement with the known literature, in addition to kras, tp53 and apc as primary events, we identify pten as a late event in the carcinogenesis, as well as nras and kras converging in igf2 amplification, the former being “selected by” tp53 mutations (`npb` 49%), the latter “selecting for” pik3ca mutations (`npb` 81%). The leftmost portion of the model links many wnt genes, in agreement with the observation that multiple concurrent lesions affecting such pathway confer selective advantage. In this respect, our model predicts multiple routes for the selection of alterations in sox9 gene, a transcription factor known to be active in colon mucosa [91]. Its mutations are directly selected by apc/ctnnb1 alterations (though with low `npb` score), by arid1a (`npb` 34%) or by fbxw7 mutations (`npb` 49%), an early mutated gene that both directly, and in a redundant way via ctnnbI, relates to sox9. The sox family of transcription factors have emerged as modulators of canonical wnt/e-catenin signaling in many disease contexts [92]. Also interestingly, fbxw7 has been previously reported to be involved in the malignant transformation from adenoma to carcinoma [93]. The rightmost part of the model involves genes from various pathways, and outlines the relation between kras and the pi3k pathway. We indeed find selection of pik3ca mutations by kras ones, as well as selection of the whole MEMO module (`npb` 64%), which is responsible for the activation of the pi3k pathway [56]. smad4 proteins relate either to kras (`npb` 34%), and fam123b (through atm) and tcf7l2 converge in dkk2 or dkk4 (`npb` 81, 17 and 34%).

*MSI-HIGH (Microsatellite Unstable)*. In agreement with the current literature, braf is the most commonly mutated gene in MSI tumors [94]. `CAPRI` predicted convergent evolution of tumors harbouring fbxw7 or apc mutations towards deletions/mutations of nras gene (`npb` 21, 28 and 54%), as well as selection of smad2 or smad4 mutations by fam123b mutations (`npb` 23 and 46%), for these tumors. Relevant to all MSI tumors seems again the role of the pi3k pathway. Indeed, a relation among apc and pik3ca mutations was inferred (`npb` 66%), consistent with recent experimental evidences pointing at a synergistic role of these mutations, which co-occurr in the majority of human colorectal cancers [95]. Similarly, we find consistently a selection trend among apc and the whole `MEMO` module (`npb` 48%). Interestingly, both mutations in apc and erbb3 select for kras mutations (`npb` 51 and 27%), which might point to interesting therapeutic implications. In contrast, mutations in braf mostly select for mutations in acvr1b (`npb` 36%), a receptor that once activated phospho-rylates smad proteins. It forms receptor complex with acvr2a, a gene mutated in these tumors that selects for tcf7l2 mutations (`npb` 34%). Tumors harbouring tp53 mutations are those selected by mutations in axin2 (`npb` 32%), a gene implicated in wnt signalling pathway, and related to unstable gastric cancer development [96]. Inactivating mutations in this gene are important, as it provides serrated adenomas with a mutator phenotype in the MSI tumorigenic pathway [97]. Thus, our results reinforce its putative role as driver gene in these tumors.

By comparing these models we can find similarity in the prediction of a potential new early event for CRC formation, fbxw7, as other authors have recently described [93]. This tumor suppressor is frequently inactivated in human cancers, yet the molecular mechanism by which it exerts its anti-tumor activity remains unexplained [98], and our models provide a new hypothesis in this respect.

## 4 Discussion

This paper represents our continued exploration of the nature of somatic evolution in cancer, and its translational exploitation through models of cancer progression, models of drug resistance (and efficacy), left - and right-censoring, sample stratification, and therapy design. Thus this paper emphasizes the engineering and dissemination of production-quality computational tools as well as validation of its applicability via use-cases carried out in collaboration with translational collaborators: e.g., colorectal cancer, analyzed jointly with epidemiologists currently studying the disease actively. As anticipated, we reasserted that the proposed model of somatic evolution in cancer not only supports the heterogeneity seen in tumor population, but also suggests a selectivity/causality relation that can be used in analyzing (epi)genomic data and exploited in therapy design - which we introduced in our earlier works [39,40]. In this paper, we have introduced an open-source pipeline, `PicNic`, which minimizes the confounding effects arising from inter-tumor heterogeneity, and we have shown that `PicNic` can be effective in extracting ensemble-level evolutionary trajectories of cancer progression.

When applied to a highly-heterogeneous cancer such as colorectal, `PicNic` was able to infer the role of many known events in colorectal cancer progression (e.g., apc, kras or tp53 in MSS tumors, and braf in MSI ones), confirming the validity of our approach^7^. Interestingly, new players in CRC progression stand out from this analysis such as fbxw7 or axin2, which deserve further investigation. In colon carcinogenesis, although each model identifies characteristic early mutations suggesting different initiation events, both models appear to converge in common pathways and functions such as wnt or mapk.

However, both models have some clear distinctive features. Specific events in MSS include mutations in intracellular genes like ctnnb1 or in pten, a well-known tumor suppressor gene. On the contrary, specific mutations in MSI tumors appear in membrane receptors such as ACvr1b, acvr2a, erbb3, lrp5, tgfbrI and tgfbr2, as well as in secreted proteins like igf2, possibly suggesting that such tumors need to disturb cell-cell and/or cell-microenvironment communication to grow. At the pathway level, genes exclusively appearing in the MSI progression model accumulate in specific pathways such as cytokine-cytokine receptor, endocytosis and tgf-*β* signaling pathway. On the other hand, genes in MSS progression model are implicated in p53, mTOR, sodium transport or inositol phosphate metabolism.

Our study also highlighted the translational relevance of the models that we can produce with `PicNic` (see Supplementary Figure S12). The evolutionary trajectories depicted by our models can, for instance, suggest previously-uncharacterized phenotypes, help in finding biomarker molecules predicting cancer progression and therapy response, explain drug resistant phenotypes and predict metastatic outcomes. The logical structure of the formulas describing alterations with equivalent fitness (i.e., the exclusivity group) can also point to novel targets of therapeutic interventions. In fact, exclusivity groups that are found to have a role in the progression can be screened for *synthetic lethality* among such genes - thus explaining why we do not observe phenotypes where such alterations co-occur. In this sense, our models describe also such clonal signatures which, though theoretically possible, are not selected. We call such conspicuously absent phenotypes *antihallmarks* [100].

Our models have other applications to both computational and cancer research. Our models, as encoded by Suppes-Bayes Causal Networks could be used as informative *generative models* for the genomic profiles for the cancer patients. In fact, as known in machine learning, such generative models are extremely useful in creating better representation of data in terms of, e.g., discriminative kernels, such as Fisher [101]. In practice, this change of representations would allow framing common classification problems in the domain of our generative structures, i.e., the models, rather than the data. As a consequence, it is possible to create a new class of more robust classification and prediction systems.

One may think of these representations as those bringing us closer to phenotypic (and causal) representation of the patient’s tumor, replacing its genotypic (and mutational) representation. We suspect that such representations will improve the accuracy of measurement of the biological clocks, dysregulated in cancer and critically needed to be measured in order to predict survival time, time to metastasis, time to evolution of drug resistance, etc. We believe that these “phenotypic clocks” can be used immediately to direct the therapeutic intervention.

Clearly, applicability and reliability of techniques such as `PicNic` is very much dependent on the background of data available. At the time of this writing, the quality, quantity and reliability of (epi)genomic data available, e.g., in public databases, is related to the ever increasing computational and technological improvements characterizing the wide area of cancer genomics. Of similar importance is the availability of wet-lab technologies for models validation. Our recent work on SubOptical Mapping technology, for instance, points to the ability to cheaply and accurately characterize translocation, indels and epigenomic modifications at the single molecule and single cell level [102,103]. This technology also provides the ability to directly validate (or refute) the hypotheses generated by `PicNic` via gene-correction and single cell perturbation approaches.

To conclude, the precision of any statistical inference technique, including `PicNic`, is influenced by the quality, availability and idiosyncrasies of the input data - the goodness of the outcomes improving along with the expected advancement in the field. Nevertheless, the strength of the proposed approach lies in the efficacy in managing possibly noisy/ biased or insufficient data, and in proposing refutable hypotheses for experimental validation.

## 5 Materials and methods

**Processing COADREAD samples with PicNic**. We instantiated `PicNic` to process clinically annotated high MSI-HIGH and MSS colorectal tumors collected from The Cancer Genome Atlas project “Human Colon and Rectal Cancer” (COADREAD) [56] – see Supplementary Figure S1. Details on the implementation and the source code to replicate this study are available as Supplementary Material. COADREAD has enough samples, especially for MSS tumors, to implement a consistent and significant statistical validation of our findings - see Supplementary Table S1.

In brief, we split subtypes by the microsatellite status of each tumor as annotated by the consortium (so, step I of `PicNic` is done by exploiting background knowledge rather than computational predictors). It should be expected that if this step is skipped or this classification is incorrect, the resulting models would noticeably differ. Once split into groups, the input COADREAD data is processed to maintain only samples for which both high-quality curated mutation and CNA data are available; for CNAs we use focal high-level amplifications and homozygous deletions.

Then, for each sample we select only alterations (mutations/CNAs) from a list of 33 driver genes manually annotated to 5 pathways in [56] - wnt, rap, tgf-*β*, pi3k and p53 (Supplementary Figures S2 and S3). This list of drivers, step II of `PicNic`, is produced by TCGA, as a result of manual curation and running MutSigCV.

In the next module of the pipeline, we fetch groups of exclusive alterations. We scanned these groups by using the MUTEX tool (Supplementary Table S2), and merged its results with the group that `TCGA` detected by using the MEMO tool, which involves mainly genes from the pi3k pathway. Knowledge on the potential exclusivity among genes in the wnt (apc,ctnnb1) and rap (kra8,nras,brap) pathways was exploited as well. Groups were then used to create `CAPRI`’s formulas; we also included hypotheses for genes which harbour mutations and homozygous deletions across different samples, see Supplementary Table S3. Data and exclusivity groups for MSS tumors are shown in Supplementary Figure S4 and S5.

CAPRI was run, as the last step of `PicNic`, on each subtype, by selecting recurrent alterations from the pool of 33 pathway genes and using both AIC/BIC regularizer. Timings to run the relevant steps of the pipeline are reported in the Supplementary Material. In the models of Figures 4 and Figure 5 each edge mirrors selective advantage among the upstream and downstream nodes, as estimated by `CAPRI`; Mann-Withney U test is carried out with statistical significance 0.05, after 100 non-parametric bootstrap iterations.

The significance of the reconstructed models and the input data is assessed by computing all the statistics/tests discussed in the Main text (temporal priority, probability raising and hypergeometric testing p-values, bootstrap and cross-validation scores). Motivation and background on each of these measures is available in the Supplementary Materials. A table with their values for edges with highest non-parametric bootstrap scores is in Supplementary Figure S8.

For the MSS cohort all the p-values are strongly significant (*p*≪0.01) except for the temporal priority of the edges connecting mutations in pam123b and atm, and erbb2 alterations (mutations and amplifications), which leads us to conclude that, even if these pairs of genes seem to undergo selective advantage, the temporal ordering of their occurrence is ambiguous and failed to be imputed correctly from the datasets, analyzed here. The same situation occurs in MSI-HIGH tumors, for the relation between kras and erbb3. Non-parametric and statistical bootstrap estimations are used to assess the strength of all the findings (Supplementary Figures S6 and S7). Moreover, any bias in the data is finally evaluated by cross-validation (Supplementary Figures S8-S11) and common statistics such as entropy loss, posterior classification and prediction errors. In general, most of the selective advantage relations depicted by the inferred models present a strong statistical support, with the MSS cohort presenting the most reliable results.

Summary implementation for COADREAD (`PicNic` steps, Figure 2): (1) TCGA clinical classification, (2) MutSigCV and TCGA manual curation, (3) MEMO, MUTEX and knowledge of wnt and RAF pathways and (4) CAPRI.

**Implement your own case study with PicNic/TRONCO**. TRONCO started as a project before `PicNic`, and is our effort at collecting, in a free R package, algorithms to infer progression models from genomic data. In its current version it offers the implementation of the `CAPRI` and CAPRESE algorithms, as well as a set of routines to pre-process genomic data. With the invention of `PicNic`, it started accommodating software routines to easily interface `CAPRI` and CAPRESE to some of the tools that we mention in Figure 2. In particular, in its current 2.0 version it supports input/output for the Matlab Network Based Stratification tool (NBS) and the Java MUTEX tool, as well as the possibility to fetch data available from the cBioPortal for Cancer Genomics (http://cbioportal.orghttp://cbioportal.org), which provides a Web resource for exploring, visualizing, and analyzing multidimensional cancer genomics data.

We plan to extend TRONCO in the future to support other similar tools and become an integral part of daily laboratory routines, thus facilitating application of `PicNic` to additional use cases.

## Authors contributions

This work follows up on our earlier project initiated by BM and carried out by Milan-Bicocca and the Catalan Institute of Oncology, based on a framework discussed at the 2014 School on Cancer, Systems and Complexity (CSAC). `PicNic` was designed and constructed by MA's Bioinformatics lab at University of Milan-Bicocca, within a project led and supervised by GC. GC, AG and DR designed the pipeline, and GC, DR and LDS coded and executed it. Data gathering and model interpretation was done by GC, LDS, DR, AG together with BM, VM and RSP. GM, MA, VM and BM provided overall organizational guidance and discussion. GC, AG, RSP and BM wrote the original draft of the paper, which all authors reviewed and revised in the final form. BM and MA are co-senior authors.

## Acknowledgments

MA, GM, GC, AG, DR acknowledge the SysBioNet project, a MIUR initiative for the Italian Roadmap of European Strategy Forum on Research Infrastructures (ESFRI) and Regione Lombardia (Italy) for the research projects RetroNet through the ASTIL Program [12-45148000-40]; U.A 053 and Network Enabled Drug Design project [ID14546A Rif SAL-7], Fondo Accordi Istituzionali 2009. BM acknowledges founding by the NSF grants CCF-0836649, CCF-0926166 and a NCI-PSOC grant. VM and RSP acknowledge the Instituto de Salud Carlos III supported by The European Regional Development Fund (ERDF) grants PI11-01439, PIE13/00022, the Spanish Association Against Cancer (AECC) Scientific Foundation, and the Catalan Government DURSI, grant 2014SGR647.

We wish to thank the anonymous reviewers for their help in improving the quality and rigor of the presentation.

## Footnotes

1 For this and other technical terms commonly used in the statistics and cancer biology communities we provide a Glossary in the Supplementary Material.

2 We mention that much attention has been recently casted on newly discovered cancer genes affecting global processes that are apparently not directly related to cancer development, such as cell signaling, chromatin and epigenomic regulation, RNA splicing, protein homeostasis, metabolism and lineage maturation [10].

3 At the time of this writing, in TCGA, sample sizes per cancer type are in the order of a few hundreds. Such numbers are expected to increase in the near future, with a clear benefit for all the statistical approaches to analyze cancer data which currently lack a proper background of data.

4 The genuine selectivity relationship sought to be inferred are subject to the vagaries of Simpson’s paradox; it can change, or worst reverse, when we try to infer them from data not suitably pre-processed. This effect (due to such paradox) manifests as data are sampled from a highly heterogenous mixture of populations of cells [40]. PiCnIc uses various mechanisms to avoid these pitfalls. In this context, it should be pointed out that input bulk sequencing data suffers also from intra-tumor heterogeneity issues, which are unfortunately intrinsic to the technology.

5 There, logical connectives such as φ (the logical “xor”) act as a proxy for hard-exclusivity, and ∨ (the logical “disjunction”) for soft one. Besides from exclusivity groups, other connectives such as logical conjunction can be used.

6 Technically, for a set of *m* alterations modeled by variables **x**_1_,…,**x**_*m*_ such a network is a Graphical Model representing the factorization of the joint distribution - ***ρ*** (***x***_1_,…,**x**_*m*_) - of observing any of the alterations in a genome (i.e., **x**_*i*_ = 1). This factorization is made compact as the model encodes the statistical dependencies in its structure via 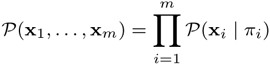 where π_*i*_ = {**x**_*j*_ | **x***j* → **x**_*i*_ ∈ ***S***̂ } are the “parents” of the *i*-th node. These are those from which the presence of the *i*-th alteration is predicted. In our approach these edges are the pictorial representation of the selective advantage relations where the alterations in π_*i*_ select for **x**_*i*_.

7 As a further investigation for CRC, we leave as future work to check whether the inferred progression are also representative of other subtyping strategies for colorectal cancer, with particular reference to recent works which show marked interconnectivity between different independent classification systems coalescing into four consensus molecular subtypes [99].

## References

[1] Nowell PC (1976) The clonal evolution of tumor cell populations. Science 194:23–23.

[2] Fidler IJ (1978) Tumor heterogeneity and the biology of cancer invasion and metastasis. Cancer Research 38:2651–2651.

[3] Dexter DL, et al. (1978) Heterogeneity of tumor cells from a single mouse mammary tumor. Cancer Research 38:3174–3174.

[4] Merlo LM, Pepper JW, Reid BJ, Maley CC (2006) Cancer as an evolutionary and ecological process. Nature Reviews Cancer 6:924–924.

[5] Hanahan D, Weinberg RA (2000) The hallmarks of cancer. CeJl 100:57–57.

[6] Hanahan D, Weinberg RA (2011) Hallmarks of cancer: the next generation. Cell 144:646–646.

[7] Huang S, Ernberg I, Kauffman S (2009) Cancer attractors: a systems view of tumors from a gene network dynamics and developmental perspective (Elsevier), No. 7, pp 869–876.

[8] Futreal PA, et al. (2004) A census of human cancer genes. Nature Reviews Cancer 4:177–177.

[9] Vogelstein B, et al. (2013) Cancer genome landscapes. Science 339:1546–1546.

[10] Garraway LA, Lander ES (2013) Lessons from the cancer genome. Cell 153:17–17.

[11] Zack TI, et al. (2013) Pan-cancer patterns of somatic copy number alteration. Nature Genetics 45:1134–1134.

[12] Baylin SB, Jones PA (2011) A decade of exploring the cancer epigenome - biological and translational implications. Nature Reviews Cancer 11:726–726.

[13] Weinberg R (2013) The Biology of Cancer (Garland Science).

[14] Albini A, Sporn MB (2007) The tumour microenvironment as a target for chemoprevention. Nature Reviews Cancer 7:139–139.

[15] Greaves M, Maley CC (2012) Clonal evolution in cancer. Nature 481:306–306.

[16] Nowak MA, Michor F, Iwasa Y (2003) The linear process of somatic evolution. Proceedings of the National Academy of Sciences 100:14966–14966.

[17] Vogelstein B, Kinzler KW (2004) Cancer genes and the pathways they control. Nature Medicine 10:789–789.

[18] Nowak MA (2006) Evolutionary Dynamics (Harvard University Press).

[19] Wood LD, et al. (2007) The genomic landscapes of human breast and colorectal cancers. Science 318:1108–1108.

[20] Jones S, et al. (2008) Core signaling pathways in human pancreatic cancers revealed by global genomic analyses. Science 321:1801–1801.

[21] Parsons DW, et al. (2008) An integrated genomic analysis of human glioblastoma multiforme. Science 321:1807–1807.

[22] Fisher R, Pusztai L, Swanton C (2013) Cancer heterogeneity: implications for targeted therapeutics. British Journal of Cancer 108:479–479.

[23] Curtis C, et al. (2012) The genomic and transcriptomic architecture of 2,000 breast tumours reveals novel subgroups. Nature 486:346–346.

[24] Gerlinger M, et al. (2012) Intratumor heterogeneity and branched evolution revealed by multiregion sequencing. The New England Journal of Medicine 366:883–883.

[25] (2015) The Cancer Genome Atlas (TCGA) https://tcga-data.nci.nih.govhttps://tcga-data.nci.nih.gov.

[26] Navin N, et al. (2011) Tumour evolution inferred by single-cell sequencing. Nature 472:90–90.

[27] Eberwine J, Sul JY, Bartfai T, Kim J (2014) The promise of single-cell sequencing. Nature Methods 11:25–25.

[28] Beerenwinkel N, Schwarz RF, Gerstung M, Markowetz F (2015) Cancer evolution: mathematical models and computational inference. Systematic biology 64:e1–e25.

[29] Ramchandani S, Bhattacharya SK, Cervoni N, Szyf M (1999) DNA methylation is a reversible biological signal. Proceedings of the National Academy of Sciences 96:6107–6107.

[30] Vogelstein B, et al. (1988) Genetic alterations during colorectal-tumor development. The New England Journal of Medicine 319:525–525.

[31] Desper R, et al. (1999) Inferring tree models for oncogenesis from comparative genome hybridization data. Journal of Computational Biology 6:37–37.

[32] Desper R, et al. (2000) Distance-based reconstruction of tree models for oncogenesis. Journal of Computational Biology 7:789–789.

[33] Szabo A, Boucher K (2002) Estimating an oncogenetic tree when false negatives and positives are present. Mathematical Biosciences 176:219–219.

[34] Beerenwinkel N, et al. (2005) Learning multiple evolutionary pathways from cross-sectional data. Journal of Computational Biology 12:584–584.

[35] Beerenwinkel N, Eriksson N, Sturmfels B (2007) Conjunctive Bayesian networks. Bernoulli pp 893–909.

[36] Gerstung M, Baudis M, Moch H, Beerenwinkel N (2009) Quantifying cancer progression with conjunctive Bayesian networks. Bioinformatics 25:2809–2809.

[37] Attolini CSO, et al. (2010) A mathematical framework to determine the temporal sequence of somatic genetic events in cancer. Proceedings of the National Academy of Sciences 107:17604–17609.

[38] Misra N, Szczurek E, Vingron M (2014) Inferring the paths of somatic evolution in cancer. Bioinformatics 17:2456?63.

[39] Olde Loohuis L, et al. (2014) Inferring tree causal models of cancer progression with probability raising. PLOS ONE 9:e115570.

[40] Ramazzotti D, et al. (2015) CAPRI: efficient inference of cancer progression models from cross-sectional data. Bioinformatics 31:3016–3016.

[41] De Sano L, et al. (2016) TRONCO: an R package for the inference of cancer progression models from heterogeneous genomic data. Bioinformatics, 10.1093/bioinformatics/ 10.1093/bioinformatics/btw035.

[42] Suppes P (1970) A Probabilistic Theory of Causality (North-Holland Publishing Company Amsterdam).

[43] Navin NE (2014) Cancer genomics: one cell at a time. Genome Biology 15:452.

[44] Wang Y, et al. (2014) Clonal evolution in breast cancer revealed by single nucleus genome sequencing. Nature 512:155–155.

[45] Gerlinger M, et al. (2014) Genomic architecture and evolution of clear cell renal cell carcinomas defined by multiregion sequencing. Nature Genetics 46:225–225.

[46] Oesper L, Mahmoody A, Raphael BJ (2013) THetA: inferring intra-tumor heterogeneity from high-throughput dna sequencing data. Genome Biol 14:R80.

[47] Oesper L, Satas G, Raphael BJ (2014) Quantifying tumor heterogeneity in whole-genome and whole-exome sequencing data. Bioinformatics 30:3532–3532.

[48] Miller CA, et al. (2014) SciClone: inferring clonal architecture and tracking the spatial and temporal patterns of tumor evolution. PLOS Computational Biology 8:e1003665.

[49] Roth A, et al. (2014) PyClone: statistical inference of clonal population structure in cancer. Nature Methods 11:396–396.

[50] Jiao W, Vembu S, Deshwar AG, Stein L, Morris Q (2014) Inferring clonal evolution of tumors from single nucleotide somatic mutations. BMC Bioinformatics 15:35.

[51] Fischer A, Vazquez-Garcia I, Illingworth CJ, Mustonen V (2014) High-definition reconstruction of clonal composition in cancer. Cell Reports 7:1740–1740.

[52] Zare H, et al. (2014) Inferring clonal composition from multiple sections of a breast cancer. PLOS Computatioanl Biology 7:e1003703.

[53] Garvin T, et al. (2015) Interactive analysis and assessment of single-cell copy-number variations. Nature methods 12:1058–1058.

[54] Malikic S, McPherson AW, Donmez N, Sahinalp CS (2015) Clonality inference in multiple tumor samples using phylogeny. Bioinformatics 31:1349–1349.

[55] El-Kebir M, Oesper L, Acheson-Field H, Raphael BJ (2015) Reconstruction of clonal trees and tumor composition from multi-sample sequencing data. Bioinformatics 31:i62–i70.

[56] The Cancer Genome Atlas Network, et al. (2012) Comprehensive molecular characterization of human colon and rectal cancer. Nature 487:330–330.

[57] Network CGAR, et al. (2013) Genomic and epigenomic landscapes of adult de novo acute myeloid leukemia. The New England Journal of Medicine 368:2059.

[58] Network CGAR, et al. (2013) Comprehensive molecular characterization of clear cell renal cell carcinoma. Nature 499:43–43.

[59] Bennett JM, et al. (1976) Proposals for the classification of the acute leukaemias french-american-british (FAB) co-operative group. British Journal of Haematology 33:451–451.

[60] Lu J, et al. (2005) MicroRNA expression profiles classify human cancers. Nature 435:834–834.

[61] Gao Y, Church G (2005) Improving molecular cancer class discovery through sparse nonnegative matrix factorization. Bioinformatics 21:3970–3970.

[62] de Souto MC, Costa IG, de Araujo DS, Ludermir TB, Schliep A (2008) Clustering cancer gene expression data: a comparative study. BMC Bioinformatics 9:497.

[63] Network CGAR, et al. (2011) Integrated genomic analyses of ovarian carcinoma. Nature 474:609–609.

[64] Konstantinopoulos PA, et al. (2010) Gene expression profile of BRCAness that correlates with responsiveness to chemotherapy and with outcome in patients with epithelial ovarian cancer. Journal of Clinical Oncology 28:3555–3555.

[65] Reis-Filho JS, Pusztai L (2011) Gene expression profiling in breast cancer: classification, prognostication, and prediction. The Lancet 378:1812–1812.

[66] Hofree M, Shen JP, Carter H, Gross A, Ideker T (2013) Network-based stratification of tumor mutations. Nature Methods 10:1108–1108.

[67] Zhong X, Yang H, Zhao S, Shyr Y, Li B (2015) Network-based stratification analysis of 13 major cancer types using mutations in panels of cancer genes. BMC Genomics 16:S7.

[68] Lawrence MS, et al. (2013) Mutational heterogeneity in cancer and the search for new cancer-associated genes. Nature 499:214–214.

[69] Gonzalez-Perez A, Lopez-Bigas N (2012) Functional impact bias reveals cancer drivers. Nucleic Acids Research 21:e169.

[70] Tamborero D, Gonzalez-Perez A, Lopez-Bigas N (2013) OncodriveCLUST: exploiting the positional clustering of somatic mutations to identify cancer genes. Bioinformatics 29:2238–2244.

[71] Dees ND, et al. (2012) MuSiC: identifying mutational significance in cancer genomes. Genome Research 22:1589–1589.

[72] Tamborero D, Lopez-Bigas N, Gonzalez-Perez A (2013) Oncodrive-CIS: a method to reveal likely driver genes based on the impact of their copy number changes on expression. PLOS ONE 8:e55489.

[73] Gundem G, et al. (2010) IntOGen: integration and data mining of multidimensional oncoge-nomic data. Nature Methods 7:92–92.

[74] Yeang CH, McCormick F, Levine A (2008) Combinatorial patterns of somatic gene mutations in cancer. The FASEB Journal 22:2605–2605.

[75] Miller CA, Settle SH, Sulman EP, Aldape KD, Milosavljevic A (2011) Discovering functional modules by identifying recurrent and mutually exclusive mutational patterns in tumors. BMC Medical Genomics 4:34.

[76] Ciriello G, Cerami E, Sander C, Schultz N (2012) Mutual exclusivity analysis identifies oncogenic network modules. Genome Research 22:398–398.

[77] Babur Ö, et al. (2015) Systematic identification of cancer driving signaling pathways based on mutual exclusivity of genomic alterations. Genome Biology 16.

[78] Vandin F, Upfal E, Raphael BJ (2012) De novo discovery of mutated driver pathways in cancer. Genome Research 22:375–375.

[79] Zhao J, Zhang S, Wu LY, Zhang XS (2012) Efficient methods for identifying mutated driver pathways in cancer. Bioinformatics 28:2940–2940.

[80] Leiserson MD, Blokh D, Sharan R, Raphael BJ (2013) Simultaneous identification of multiple driver pathways in cancer. PLOS Computational Biology 5:e1003054.

[81] Leiserson MDM, Wu HT, Vandin F, Raphael BJ (2015) CoMEt: a statistical approach to identify combinations of mutually exclusive alterations in cancer. Genome Biology 16:160.

[82] Hua X, et al. (2015) MEGSA: A powerful and flexible framework for analyzing mutual exclusivity of tumor mutations. bioRxiv http://dx.doi.org/10.1101/027474 http://dx.doi.org/10.1101/027474.

[83] Szczurek E, Beerenwinkel N (2014) Modeling mutual exclusivity of cancer mutations. PLoS Computational Biology 10.

[84] Leiserson MD, et al. (2015) Pan-cancer network analysis identifies combinations of rare somatic mutations across pathways and protein complexes. Nature Genetics 47:106–106.

[85] Vandin F, Upfal E, Raphael BJ (2011) Algorithms for detecting significantly mutated pathways in cancer. Journal of Computational Biology 18:507–507.

[86] Efron B, Tibshirani RJ (1994) An Introduction to the Bootstrap (CRC press).

[87] Koller D, Friedman N (2009) Probabilistic Graphical Models: Principles and Techniques - Adaptive Computation and Machine Learning (The MIT Press).

[88] Ogino S, Goel A (2008) Molecular classification and correlates in colorectal cancer. The Journal of Molecular Diagnostics 10:13–13.

[89] Fearon ER, Vogelstein B (1990) A genetic model for colorectal tumorigenesis. Cell 61:759–759.

[90] Vilar E, Gruber SB (2010) Microsatellite instability in colorectal cancer - the stable evidence. Nature reviews Clinical oncology 7:153–153.

[91] Abdel-Samad R, et al. (2011) MiniSOX9, a dominant-negative variant in colon cancer cells. Oncogene 30:2493–2493.

[92] Kormish JD, Sinner D, Zorn AM (2010) Interactions between sox factors and wnt/*β*-catenin signaling in development and disease. Developmental Dynamics 239:56–56.

[93] Li L, et al. (2014) Sequential expression of miR-182 and miR-503 cooperatively targets FBXW7, contributing to the malignant transformation of colon adenoma to adenocarcinoma. The Journal of Pathology 234:488–488.

[94] Kim JH, Kang GH (2014) Molecular and prognostic heterogeneity of microsatellite-unstable colorectal cancer. World Journal of Gastroenterology 20:4230.

[95] Deming DA, et al. (2014) PIK3CA and APC mutations are synergistic in the development of intestinal cancers. Oncogene 33:2245–2245.

[96] Kim MS, Kim SS, Ahn CH, Yoo NJ, Lee SH (2009) Frameshift mutations of Wnt pathway genes AXIN2 and TCF7L2 in gastric carcinomas with high microsatellite instability. Human Pathology 40:58–58.

[97] Muto Y, et al. (2014) DNA methylation alterations of AXIN2 in serrated adenomas and colon carcinomas with microsatellite instability. BMC Cancer 14:466.

[98] Zhan P, et al. (2015) FBXW7 negatively regulates ENO1 expression and function in colorectal cancer. Laboratory Investigation 9:995?1004.

[99] Guinney J, et al. (2015) The consensus molecular subtypes of colorectal cancer. Nature medicine, in print.

[100] Loohuis LO, Witzel A, Mishra B (2014) Cancer hybrid automata: model, beliefs and therapy. Information and Computation 236:68–68.

[101] Korsunsky I (2016) Ph.D. thesis (New York University).

[102] Reed J, et al. (2012) Identifying individual dna species in a complex mixture by precisely measuring the spacing between nicking restriction enzymes with atomic force microscope. Journal of The Royal Society Interface 9:2341–2341.

[103] Sundstrom A, et al. (2012) Image analysis and length estimation of biomolecules using afm. IEEE Transactions on Information Technology in Biomedicine 16:1200–1200.

